# Building an Interoperable Rare Disease Multi-omic Resource: The GREGoR Data Model and Dataset

**DOI:** 10.64898/2026.05.15.725546

**Authors:** B D Heavner, M M Wheeler, J D Bengtsson, C M Carvalho, W A Cheung, M P Conomos, E C Délot, S DiTroia, V S Ganesh, S M Gogarten, C M Grochowski, S N Jhangiani, C H King, C LeMaster, C T Marvin, S Marwaha, D E Miller, A O’Donnell-Luria, L Pais, K Patterson, G Qi, M Richardson, C Smail, A M Stilp, C C Tong, R A Ungar, B Weisburd, M J Bamshad, J A Bernstein, E E Eichler, R A Gibbs, J R Lupski, S May, S B Montgomery, T Pastinen, J Posey, H L Rehm, A Shojaie, M E Talkowski, E Vilain, C Wei, M T Wheeler, Q Yi, Genomics Research to Elucidate the Genetics of Rare Diseases (GREGoR) Consortium, GREGoR Consortium Data Standards and Analysis Working Group, S I Berger, J X Chong

## Abstract

Rare disease research and diagnosis rely on the integration of genomic and phenotypic data generated across diverse clinical sites; however, the absence of widely adopted standards for representing genomic data and associated metadata has limited data interoperability, reuse, and cross-study analysis. The Genomics Research to Elucidate the Genetics of Rare Diseases (GREGoR) Consortium was established to investigate challenging rare disease cases and evaluate emerging multi-omic technologies for clinical translation. To support coordinated data integration across distributed research sites, we developed a common Consortium Data Model in partnership with domain experts to standardize the capture of participant-, family-, phenotype- and assay-level metadata, with a particular emphasis on using a modular architecture to support linking of multiple data versions from multiple omic technologies to a single individual and attribution of a genetic finding to the specific technology used for its initial discovery. Adoption of the GREGoR Data Model has enabled continued generation and public release of a harmonized, analysis-ready Consortium Dataset. The most recent release includes phenotypic, family and multi-omic data from 12,292 participants in 5,029 families. Other rare disease data sharing efforts are beginning to adopt this data model which will facilitate cross consortium analyses and empower rare disease research. This work demonstrates that a collaborative, flexible, and scalable data model can enable large-scale rare disease research, facilitate cross-center data harmonization, and enable data interoperability.

## Introduction

Rare and undiagnosed disease research presents persistent challenges for both discovery and diagnosis. Delineation of new rare diseases and efforts to characterize the spectrum of clinical findings associated with a rare disease has relied on the collection and comparison of individual cases with overlapping features. However, individuals with rare disease are dispersed globally, making it difficult for individual researchers to reach multiple families with the same rare condition^1^. It can be impractical for individual investigators to assemble sufficiently large and deeply phenotyped cohorts to support discovery. As the field advances efforts to resolve challenging “exome-negative” cases^1,2^, data sharing has become essential for enabling gene discovery, improving variant interpretation, and supporting clinical translation^3^.

The Genomics Research to Elucidate the Genetics of Rare Diseases (GREGoR) Consortium was launched to study thousands of challenging rare disease cases and families while applying, evaluating, and standardizing approaches using emerging genomics technologies^4^. GREGoR is a collaboration of five Research Centers which contribute participant, family, phenotype and multi-omic experimental data. These data are submitted to the GREGoR Data Coordinating Center (DCC) for harmonization, quality control, and dissemination as a coherent resource for the broader research community.

GREGoR has been generating data within a broader landscape of rapidly evolving data-sharing and deposition policies, increased security requirements for NIH-funded research^5^, and the establishment of cloud-based NIH data repositories such as the NHGRI’s Analysis Visualization and Informatics Lab-space (AnVIL)^6^. The shift from on-premises computation to cloud-based environments enhances data security and decreases the need for researchers to maintain extensive local infrastructure. However, it also introduces new challenges including the need for coordinated data governance and standardized data structures to support shared and scalable analyses.

One of the strengths of the GREGoR dataset is the breadth and scale of multi-omic and phenotypic data collected from rare disease patients and their families. Joint analysis of harmonized datasets across research centers has the potential to increase diagnostic yield and accelerate disease–gene discovery beyond what could be achieved through isolated analyses^3,7^. However, this requires standardized metadata and clearly defined data structures to enable data integration across technologies, studies, and analytic frameworks^8^. Without such standards, differences in experimental protocols, sequencing platforms, and phenotypic annotation can limit interoperability, complicate cross-study analyses, and introduce technical confounding. To address these challenges, we developed the GREGoR Data Model – a structured framework that defines the metadata requirements for participant, family, phenotypic and multi-omic data for rare disease research. The model was designed to support interoperability, facilitate data validation and submission, enable iterative data release, and promote secondary use of consortium data. By establishing shared conventions for representing complex datasets, the GREGoR Data Model provides a foundation for scalable rare disease research and advances the development of a more unified and reusable data ecosystem.

## Materials and Methods

### Design Principles

GREGoR data are collected from five Research Centers, each contributing participant, family, phenotype, and diverse multi-omic data. These data are generated through a variety of experimental pipelines and assays and are submitted to the GREGoR DCC. Together, the Research Centers and the DCC defined a shared consortium data model based on the following principles:

#### Research utility and adaptability

we developed a flexible data model designed to advance rare disease research while remaining adaptable for unforeseen future applications. To ensure long-term utility, data must be accompanied by sufficient context—such as versions of bioinformatic tools, details of wet-lab procedures and handling, and phenotypic information from family members in extended pedigrees without -omic data available that still may influence analysis. In designing the model, we applied a simple heuristic: subject matter experts considered what information they would need to meaningfully analyze a given data type, or conversely, what metadata they would provide to others for the same purpose.

#### Interoperability

we designed the GREGoR Data Model to encourage the use of structured vocabularies or ontologies. Previous experience demonstrates that using structured vocabularies or defined ontologies facilitates combining data across studies or transforming data for cross-study harmonization^9^. At the same time, we recognize that there are not defined vocabularies or ontologies for all concepts that are necessary for working with rare disease data. Therefore, in some cases, the GREGoR Data Model specifies and restricts a field to a predefined set of allowable values. Finally, recognizing that there are also circumstances where a more general description of a metadata feature is required, the Data model also includes some free text fields for cases where defined ontologies or enumerated value approaches are not feasible or sufficient.

#### Ease of use

the data model is designed to avoid requiring programming expertise for data submission. This is achieved by instantiating the data model as a series of linked tables that can be edited in a spreadsheet program. However, the data model can also be serialized in formats that are amenable to programmatic interaction, such as JSON, LinkML, or DBML. Rather than being prescriptive about the encoding of the fields defined in the GREGoR Data Model, we emphasize tools to convert among these formats. We regard the JSON version of the data model as the “canonical” version and track it in a version-controlled GitHub repository (https://github.com/UW-GAC/gregor_data_models)^10^, which also contains the code for converting between alternate formats.

#### Flexibility and modularity

we designed the Data Model to accommodate the use of many emerging technologies and novel data types for rare disease research. The GREGoR Consortium includes domain experts who specialize in a variety of molecular data types, including short-read DNA sequencing, long-read DNA sequencing, short-read RNA-seq, and short-read ATAC-seq. Development of the GREGoR Data Model involved active, interdisciplinary collaboration with subject matter experts to support various data types. This structured approach to data sharing provides a mechanism for the Data Model to be expanded to include additional data that researchers may want to contribute to the GREGoR Dataset in the future. This modular approach provides the Data Model with the flexibility to evolve over time as bioinformatic and experimental protocols mature and best practices are refined.

#### Person-centric

the operable unit of the GREGoR Data Model is a research participant - a proband or relative from whom information and/or biospecimens have been obtained. This design explicitly links participants to families, phenotypes, biospecimens, experiments, and downstream analytic outputs, enabling consistent representation of inheritance, solve status, and multi-omic context.

The GREGoR Data Model is designed to organize and standardize metadata rather than to function as a variant warehouse or phenotype-matching engine. For example, variant files (e.g., VCFs) are referenced through structured metadata fields but are not stored as queryable variant-level databases within the model itself. Similarly, phenotype data are represented in computable form intended to interoperate with external analytic platforms rather than replace dedicated case-matching or variant interpretation systems.

#### Transparency and documentation

to facilitate interoperability of the Data Model itself, we implemented a collaborative process to produce versioned releases and encoding the GREGoR Data Model in a way that can be machine-parsable which facilitates future data transformations. Although the GREGoR Consortium leverages the computational infrastructure of the AnVIL platform, the GREGoR Data Model is translatable and platform-independent.

### Organization of the GREGoR Data Model

The GREGoR Data Model is a collection of data tables with defined relationships that capture the structure and interconnections among key entities. Following the design principles listed above, the tables are based on a conceptual hierarchy with core, required tables linking participants to families, available biospecimens, and optional tables describing -omics and other experimental data and subsequent bioinformatic analyses (**Figure 1A, Document S1-S2**). Optional components include experiment, aligned, called variant tables, each specific to a given assay or -omics data type as well as a genetic finding table that captures interpreted candidate variant information.

**Figure 1.**
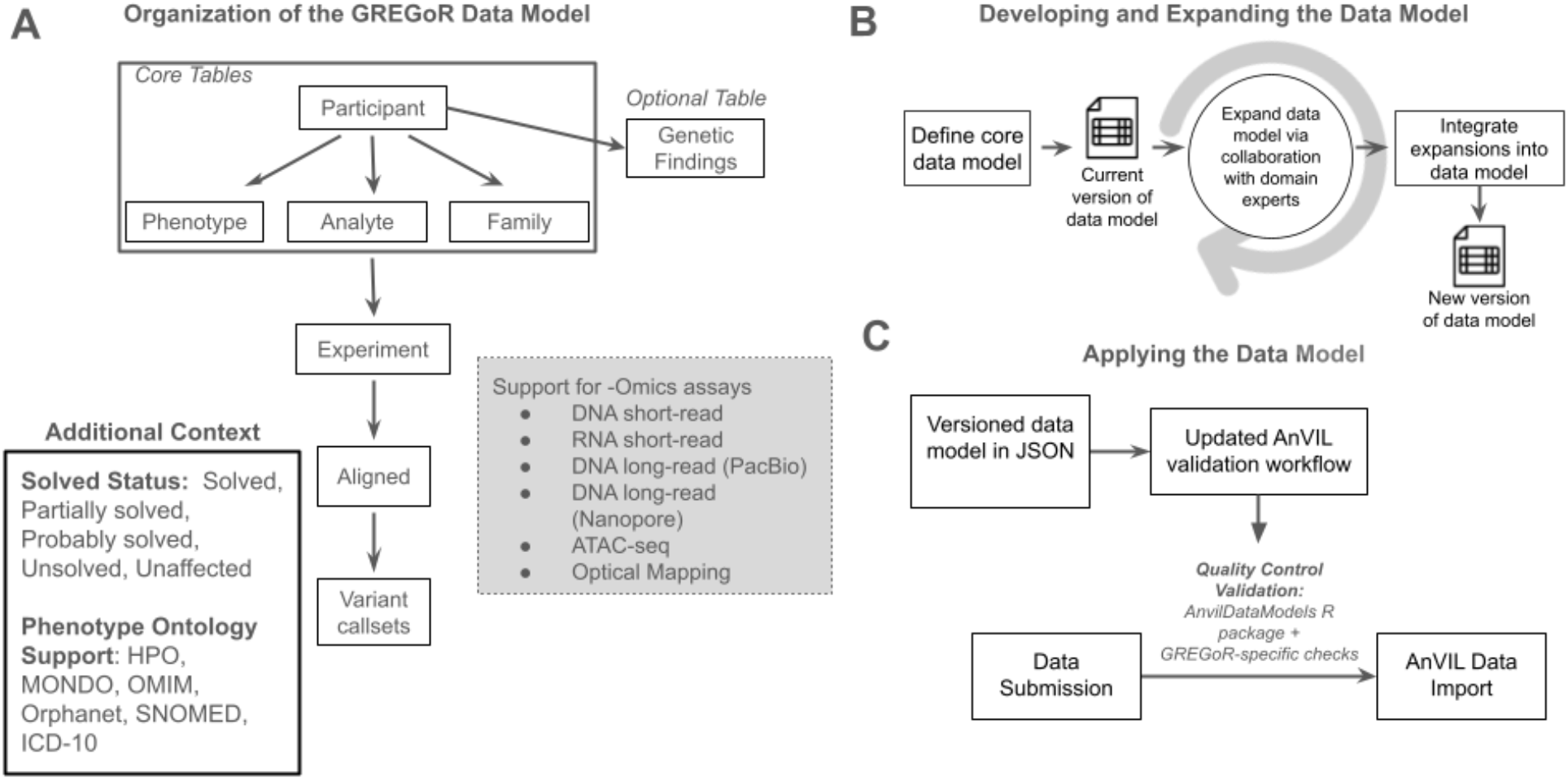
Organization, development and implementation of the GREGoR Data Model. (A) Organization of the GREGoR Data Model. The GREGoR Data Model is centered on a Participant table that links core tables capturing Phenotype, Analyte, and Family metadata. Experimental data are structured through Experiment tables that connect to aligned files and derived variant callsets. The model supports multiple -omics assays, including DNA short-read, DNA long-read (PacBio or Nanopore), optical genome mapping, RNA short-read, RNA long-read, and ATAC-seq. A Genetic Findings table integrates discoveries across these technologies, capturing curated candidate variants. The framework also incorporates standardized solve status classifications and structured phenotype ontology to enable consistent representation (see Supplementary Information for additional detail). **(B) Developing and expanding the GREGoR Data Model**. The GREGoR Data Model was developed through an iterative, versioned process in which a core schema is defined and subsequently expanded through collaboration with domain experts. Proposed extensions are evaluated, integrated into the schema, and released as new versions, enabling the model to accommodate additional assay types, metadata standards, and analytical requirements. **(C) Applying the data model**. The versioned data model is implemented in JSON and operationalized through AnVIL-based validation workflows. Data submissions undergo automated quality control using the AnvilDataModels R package together with GREGoR-specific validation checks prior to import into AnVIL.

The GREGoR Data Model currently consists of 38 linked tables (Version 1.11, https://github.com/UW-GAC/gregor_data_models, **Document S1-S2**), including 4 core tables required for quality control and workflow-based import into AnVIL and 34 optional tables for experimental metadata and genetic findings. Within each table, there are mandatory elements, as well as optional fields to provide additional context and information. This approach, which combines required and optional fields, enables comprehensive metadata capture while preserving flexibility.

The current version of the GREGoR Data Model applies a modular approach to support data for the following experiment types: DNA short-read, DNA long-read (PacBio or Nanopore), optical genome mapping, RNA short-read, RNA long-read, and ATAC-seq (**Figure 1A**). The GREGoR Data Model is implemented on the NHGRI AnVIL cloud platform^6^, with minor adaptations to accommodate platform-specific table formats. For example, the concept of a “set” of tables linking collections of experiments to jointly called VCF files comes from AnVIL’s built-in support for set tables.

### Design features that support rare disease analyses

To illustrate the key design features of the GREGoR Data Model, we use the participant and phenotype-level representation as examples to highlight analytically motivated design choices that support rare disease analysis (detailed field-level definitions for the full data model are provided as supplemental materials in **Document S1**).

The participant table includes demographic information such as sex, race and ethnicity, ancestry, and age at enrollment. It also includes information about the affected status, primary phenotype, and solve status for participants. The ‘sex’ field is defined as “biological sex assigned at birth,” with the enumerated values of “Male”, “Female”, or “Unknown”, following the conventions used by the All of Us Research Study^11^ which adopted these values from the Observational Medical Outcomes Partnership (OMOP) common data model^12^. The GREGoR Data Model includes an optional free-text, ‘sex-detail’ field that can record known discrepancies between the expected sex chromosome (i.e., female=>XX, male=>XY) and the actual karyotype as well as known differences of sex development (DSD).

The participant table also includes ‘reported_race’, ‘reported_ethnicity’, and ‘ancestry_detail’ fields to support self-identified and genetic ancestry descriptions. Reported_race and reported_ethnicity fields follow US OMB guidelines, with enumerated values including “American Indian or Alaska Native,” “Asian,” “Black or African American,” “Native Hawaiian or Other Pacific Islander,” “Middle Eastern or North African” and “White” for race, and “Hispanic or Latino” or “Not Hispanic or Latino” for ethnicity. The free-text ancestry_detail field allows further description of participant ancestry (e.g., Alaska Native, Ashkenazi Jewish, Egyptian, German) or be used for genetically-inferred ancestry.

The affected_status field indicates whether a participant is affected with the primary phenotype under investigation in the family. The solve_status field reflects whether the submitting GREGoR Research Center considers the primary phenotype to be genetically explained (“solved”). Enumerated values for solve_status include: Solved, Partially solved, Probably solved, Unsolved, and Unaffected.

The GREGoR Consortium defined these categories to promote consistency across Research Centers, acknowledging that criteria for determining a case as solved may vary depending on evidence strength, gene–disease validity, and available functional data. Brief definitions for GREGoR solve_status categories are shown in **Table 1**, with full definitions stratified by mode of inheritance are described in the supplementary materials **(Table S1-S2)**. Solve status definitions were informed by the ClinGen gene–disease validity framework, ACMG/AMP sequence variant classification guidelines, and ACMG/ClinGen CNV classification standards^13–15^.

**Table 1.**
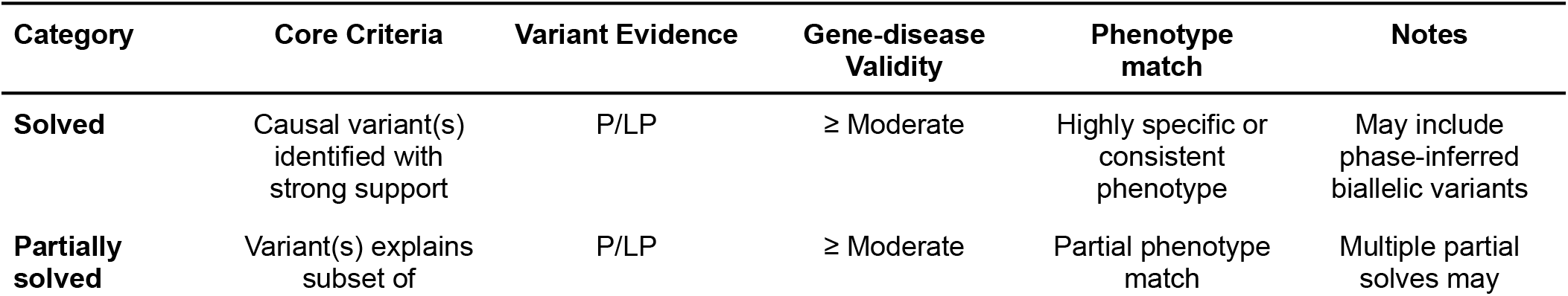
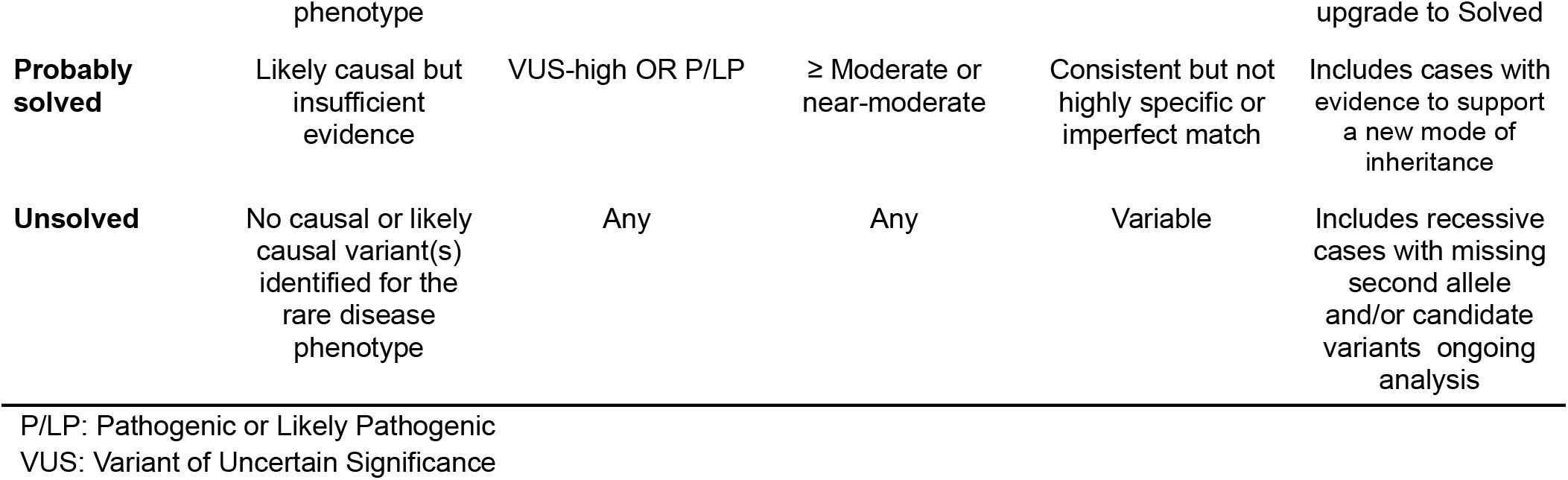
Criteria for Solve Status Classification.

| Category | Core Criteria | Variant Evidence | Gene-disease Validity | Phenotype match | Notes |
| --- | --- | --- | --- | --- | --- |
| **Solved** | Causal variant(s) identified with strong support | P/LP | ≥ Moderate | Highly specific or consistent phenotype | May include phase-inferred biallelic variants |
| **Partially solved** | Variant(s) explains subset of | P/LP | ≥ Moderate | Partial phenotype match | Multiple partial solves may |

|  | phenotype |  |  |  | upgrade to Solved |
| --- | --- | --- | --- | --- | --- |
| **Probably solved** | Likely causal but insufficient evidence | VUS-high OR P/LP | ≥ Moderate or near-moderate | Consistent but not highly specific or imperfect match | Includes cases with evidence to support a new mode of inheritance |
| **Unsolved** | No causal or likely causal variant(s) identified for the rare disease phenotype | Any | Any | Variable | Includes recessive cases with missing second allele and/or candidate variants ongoing analysis |

P/LP: Pathogenic or Likely Pathogenic
VUS: Variant of Uncertain Significance

The genetic findings table links reported variants to participants and integrates these data with family structure and phenotype information, enabling systematic interpretation of variant–phenotype relationships across the cohort.

The phenotype table supports entry of list of term identifiers from standardized phenotypic ontologies (HPO, MONDO, OMIM, Orphanet, SNOMED, and ICD-10) to facilitate annotation of the presence or absence of specific phenotypes. Free-text fields accommodate additional details. Each participant may have multiple phenotype entries, enabling representation of complex or detailed phenotypic profiles. Affected participants are required to have at least one phenotype recorded.

### Implementation of the Data Model

Implementation of the GREGoR Data Model required cross-consortium collaboration for defining the tables and features necessary to enable collaborative primary and secondary analysis, along with supporting technical infrastructure to encode, submit, and share metadata tables (**Figure 1B-C**). We used an asynchronous framework consisting of shared documents to collect input and record comments that could be resolved via the Data Standards and Analysis Working Group, followed by an iterative, version-controlled release process implemented by the DCC to progressively refine the model over the course of the Consortium (**Figure 1B**).

The process of making the GREGoR Data Model began by defining the overall hierarchy and conceptual organization of the data, then establishing a “core” minimal model that encompasses essential fields and tables. In addition to being a practical starting point for expanding the data model to new data types, this minimal core was also considered useful to ensure broad applicability across disease types and facilitate community adoption. After the core model was defined, initial expansion focused on adding optional tables to describe mature, widely adopted experimental technologies, starting with short-read DNA and RNA sequencing. Subject matter experts proposed domain-specific expansions, which were reviewed, refined, and integrated. Expansion and refinement proceeded through standardized version updates, with regular evaluation and versioned releases via GitHub. Over time, the GREGoR Data Model has added optional tables to expand coverage to newer experimental technologies, such as long-read sequencing, optical genome mapping, and other data types.

Aligning with our Design Principles, we prioritized implementing the GREGoR Data Model in a format that would minimize barriers for contributing Research Centers and Partner Members. Specifically, the metadata fields defined by the Data Model are implemented as columns in a columnar data format, such as a .tsv or .csv file or spreadsheet. Using the metadata and format defined by the Data Model, data submitters used AnVIL workspaces for collaborative data sharing (**Figure 1C**). Quality control and validation workflows ensured that data tables conformed to the model during the data submission process, as the tabular data was imported to AnVIL Workspace Data Tables. The validation workflows ensured that required fields were populated with allowable values and relationships between tables were maintained across submission cycles.

For validation, the GREGoR Data Coordinating Center developed the AnvilDataModels R package (https://github.com/UW-GAC/AnvilDataModels), which reads a JSON-formatted data model and verifies consistency of submitted tables. This package is generalizable and can be used with other data models. A complementary workflow description language (WDL) runs functions from this package within an AnVIL compute instance. The workflow generates a validation report, which describes any discrepancies between the submitted data and the data model. If there are no discrepancies and the validation is successful, the workflow imports the data into AnVIL tables (https://dockstore.org/workflows/github.com/UW-GAC/anvil-util-workflows/validate_data_model: main). Additionally, a GREGoR-specific workflow performs model-specific checks, such as verifying that sample identifiers in file headers match table entries and that phenotype ontology terms are correctly formatted (https://dockstore.org/workflows/github.com/UW-GAC/gregor-file-checks/validate_gregor_model :main).

### The GREGoR Data Model and AnVIL

The GREGoR Data Model is designed to be portable across diverse computational environments. Conformant datasets can be analyzed using on-premises or cloud-based infrastructures. Here, the GREGoR Consortium has partnered closely with the AnVIL platform, which provides features that support data analysis formatted according to the GREGoR Data Model.

Within AnVIL workspaces, the tables of the GREGoR Data Model are implemented as *Data Tables*, a key workspace feature. These tables are searchable using the AnVIL Web Interface, or programmatically via Python or R scripts^16^. Although AnVIL data tables do not constitute a relational database, they are structured to enable SQL-like joins across related entities, facilitating flexible cohort selection and multi-table queries.

More broadly, the GREGoR Data Model includes all of the necessary metadata elements required for indexing within AnVIL’s search infrastructure as a NIH-designated data repository. By conforming to the GREGoR Data Model, data submitted to AnVIL for controlled-access release can be programmatically ingested into the AnVIL Data Catalogue – improving the findability, interoperability and reuse of submitted data.

Furthermore, the GREGoR Data Model has been adopted by AnVIL as an example consortium-level data model and is made available to researchers preparing their data for archival and controlled-access release.

## Results

### The GREGoR Dataset

We implemented the GREGoR Data Model to organize and standardize metadata across the Consortium Dataset. Adherence to the data model ensured completeness and standardization of all required fields. Optional field completion varied substantially across tables. Experiment-level tables exhibited relatively high and consistent completion (e.g., median completeness >0.5 in several assay-specific tables), indicating that many optional metadata fields are routinely captured. In contrast, the analyte and family tables showed minimal optional field usage (median ≈ 0), suggesting that most optional attributes in these tables are not routinely collected. Intermediate patterns were observed for participant and phenotype tables, with evidence of uneven usage across optional fields **(see Table S3)**.

Implementation of the data model enabled systematic evaluation of the GREGoR Dataset and facilitated rapid controlled data release cycles. Key statistics describing cohort composition, phenotypic diversity, and multi-omic data availability across public releases are shown in **Figure 2A-B**.The current release of the GREGoR Dataset includes 12,292 participants from 5,029 families suspected to have a rare disease, most of whom lacked an explanatory genetic variant(s) following prior clinical or research genetic testing (**Figure 2A**), most commonly exome sequencing. The cohort is predominantly composed of trios (N= 2,367, 47%), followed by singletons (N=1436, 28.6%), duos (N=675, 13.4%), and larger family structures (N=551, 11%). At present, 84.1% of probands (N = 4,130) remain undiagnosed, underscoring the focus of GREGoR on challenging rare disease cases.

**Figure 2.**
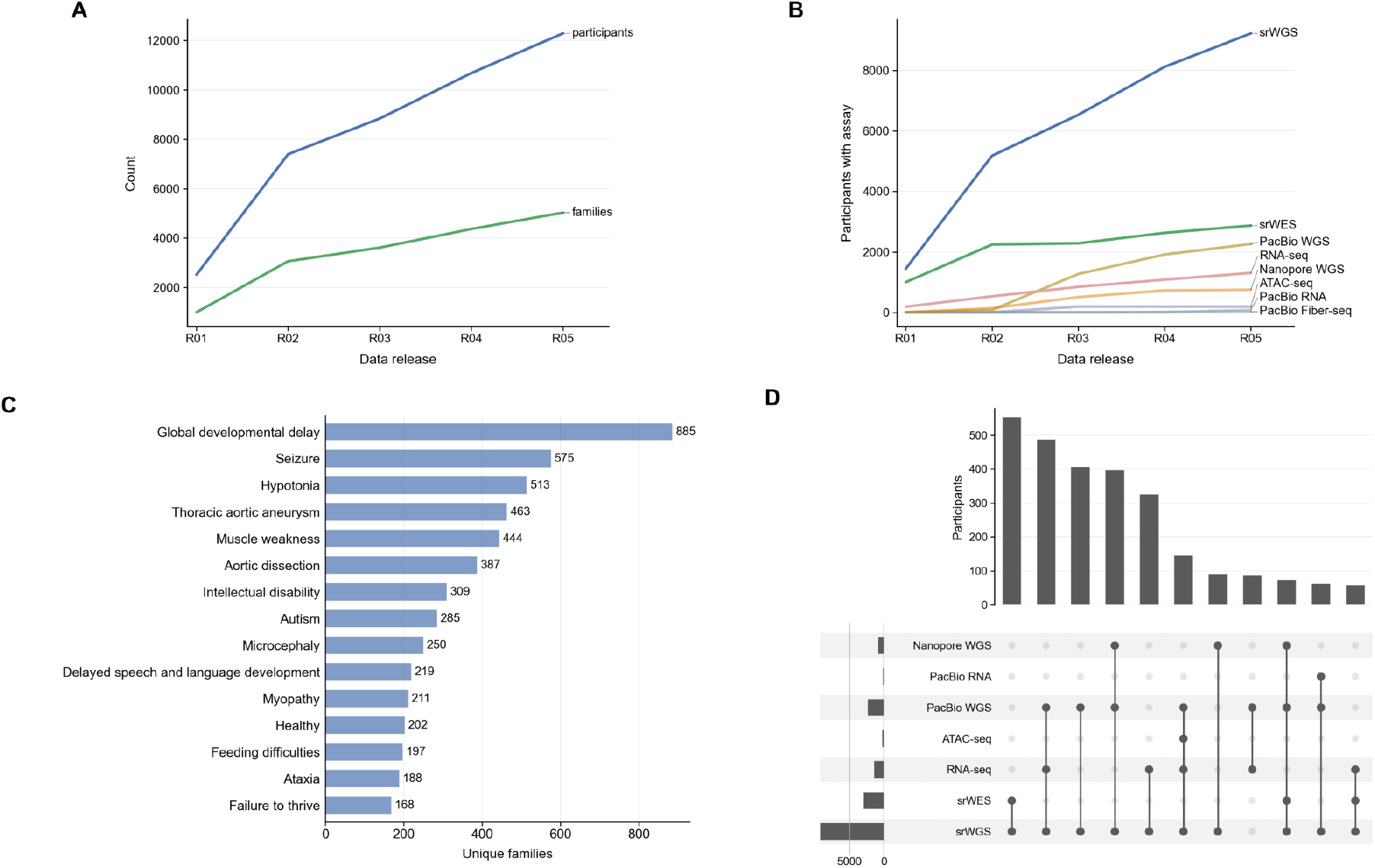
Overview of the GREGoR Dataset. (A) Growth of the cohort across controlled-access data releases distributed via dbGaP and the NHGRI AnVIL platform, showing the number of participants and families included in each release. (B) Expansion of assay types across data releases, including short-read whole-genome sequencing (srWGS), short-read exome sequencing (srWES), short-read RNA sequencing (RNA-seq), long-read whole-genome sequencing (lrWGS; PacBio and Oxford Nanopore), short-read ATAC-seq, long-read RNA sequencing (PacBio), and long-read Fiber-seq (PacBio). (C) Most prevalent Human Phenotype Ontology (HPO) terms across unique families in the current data release, shown in descending order of frequency. (D) Overlap of assay types among participants in the current data release, shown in descending order of intersection size using an UpSet plot. Only intersections involving more than one assay and comprising more than 50 participants are shown.

Participant metadata indicate that 53.9% of probands (N = 2,650) are male and 45.4% (N = 2,229) are female, while 0.7% (N = 34) are recorded as “unknown” for biological sex at birth. Racial descriptors were reported by investigators when available and are present for approximately 73% of probands. Among those with reported race data, most identify as White (78.6%), followed by Asian (8.5%), Middle Eastern (6.8%), and African or African American (4.8%), with less than 1% identifying as Native American or Alaska Native or Native Hawaiian or Other Pacific Islander.

Affected status information is available for all participants (probands = 40%, affected family member = 6.7%, possibly affected family member = 0.4%, unaffected participants = 49.9 %, unknown = 3.1%). Standardized phenotypic ontology terms are provided for nearly all probands and other affected individuals (probands = 98.8%, other affected = 100%, possibly affected = 80%). The dataset contains a total of 3,450 unique phenotypic ontology terms (HPO = 3,375; MONDO = 8; OMIM = 32; Orphanet = 35), with an average of 5.8 HPO terms per proband (range: 1–68). Analysis of HPO term distributions across families indicates a heterogeneous rare disease cohort, with neurodevelopmental, neuromuscular, and syndromic conditions most frequently represented (**Figure 2C**).

### Multi-omic Data

Across the cohort, short-read genome or exome sequencing data are available for the majority of participants (N = 11,358; **Figure 2B**). A subset of short-read genome data was harmonized by the GREGoR Data Coordinating Center and used to generate a consortium-wide joint callset that includes variant calls (single-nucleotide variants [SNVs] and small insertions/deletions [indels]) generated with GATK-DRAGEN for 3,397 individuals. Additional multi-omic datasets include short-read RNA-seq (N = 1,303) and ATAC-seq (N = 186), as well as long-read sequencing (PacBio genome = 2,261; PacBio RNA-seq = 85; PacBio FiberSeq = 24; Oxford Nanopore genome = 748) (**Figure 2B**). A substantial subset of participants has overlapping multi-omic data, enabling integrative analyses across sequencing modalities (**Figure 2D**). Notably, 209 probands have matched short-read genome sequencing, long-read PacBio genome sequencing, and short-read RNA-seq data. An additional 164 probands have both short-read exome and short-read genome data, while 144 probands have matched short-read genome, long-read PacBio genome, and long-read Nanopore genome data.

### Candidate Genetic Findings

GREGoR Research Centers have contributed candidate genetic findings with varying levels of supporting evidence, providing a structured view of prioritized variants across the cohort. Within the GREGoR Data Model, each genetic finding is linked to a specific participant, their associated phenotype annotations, and participant-level solve status, enabling integrated representation of variant, individual, and clinical interpretation data.

In the current GREGoR release (R05), the candidate genetic findings table contains 2,265 reported findings for 1,615 participants, representing a subset of the 12,292 participants in the overall release. For the analyses and supplementary tables presented here, we performed additional quality control and harmonization, including review of variant representation, transcript and HGVS annotation, and manual curation of potential duplicate entries. Findings that could not be confidently reconciled during QC were removed from the manuscript version of the table. As a result, the publication table (**Table S4**) represents a curated subset of the original R05 candidate findings release, comprising 2,252 findings from 1,606 unique participants.

Genetic findings in **Table S4** span a range of solve status categories, reflecting varying levels of diagnostic resolution across the cohort. Most findings are associated with unsolved cases (N = 1,205), followed by solved (N = 803), probably solved (N = 198), and partially solved (N = 37). In **Figure 3**, we show solve status collapsed to the participant level (N = 1,606). This reduces counts across all categories but preserves the overall distribution, underscoring both the progress made in identifying likely causal variants and the substantial proportion of cases that remain unresolved (including cases with and without candidates of interest).

**Figure 3.**
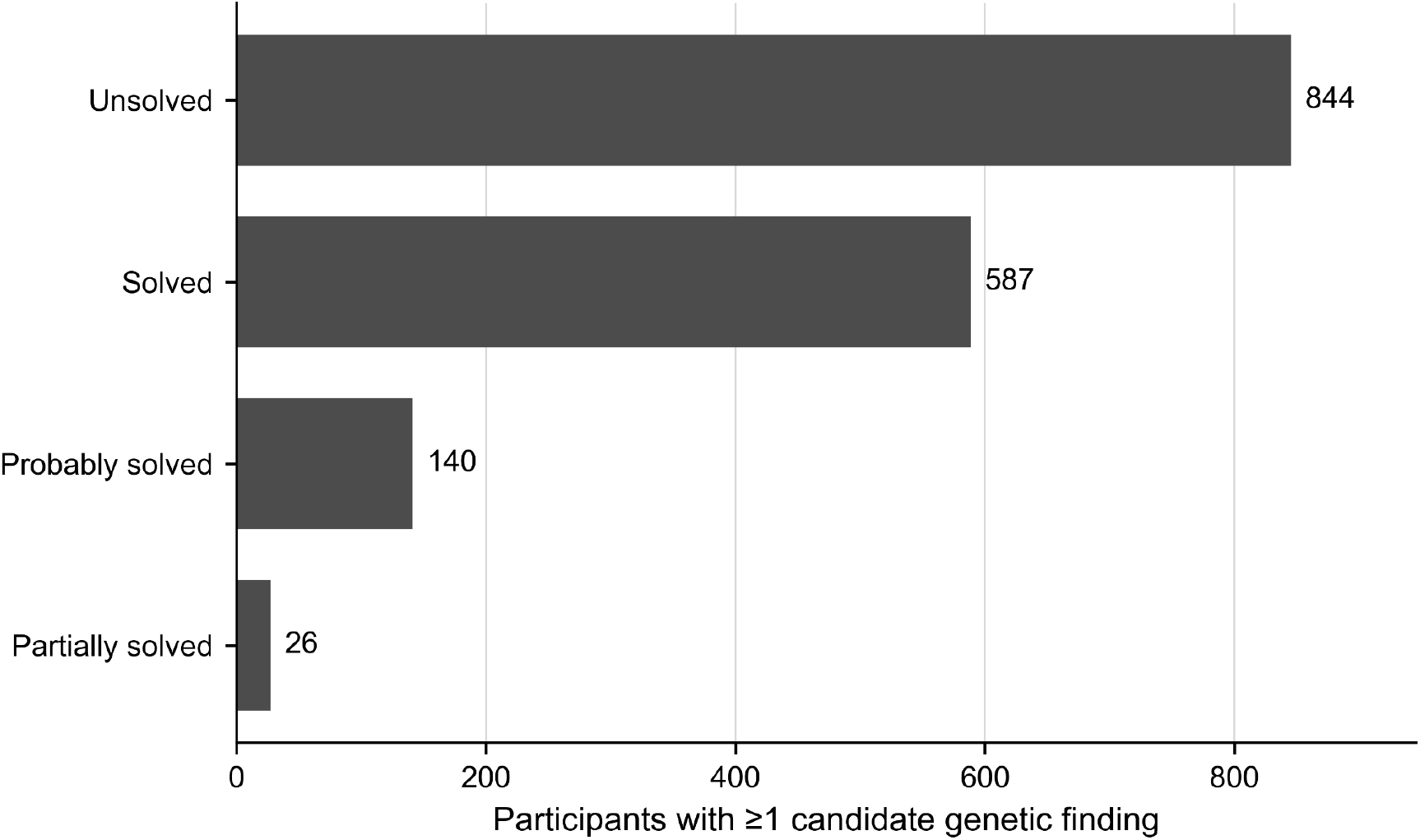
Solve status across participants with candidate genetic findings. Participants with genetic findings are categorized according to GREGoR solve status definitions. Most participants with a reported finding remain unsolved (N = 844), followed by solved (N = 587), probably solved (N = 140), partially solved (N = 26), highlighting both progress in variant interpretation and the substantial proportion of cases that remain unresolved.

**Table S4** includes genetic findings for SNVs (N = 1,722), indels (N = 350), structural variants (N = 156), and repeat expansions or contractions (N = 25), collectively involving a broad set of known disease genes (unique genes = 641), as well as candidate novel disease genes and potential phenotypic expansions (unique genes = 688). Genetic findings associated with solved or probably solved cases (N = 1,001) include SNVs (N = 708), indels (N = 183), structural variants (N = 95), and repeat expansions or contractions (N = 15), and collectively involve known disease genes (unique genes = 442) as well as candidate novel disease genes or potential phenotypic expansions (unique genes = 146).

Most candidate findings were identified using short-read exome or genome sequencing (94.4%), with a smaller subset identified using long-read genome sequencing, either alone or in combination with other technologies (3.5%). A small subset of findings were detected using short-read or long-read RNA-seq (1.2%) or via prior (pre-GREGoR) clinical gene panel sequencing or SNP microarrays (1.1%). Together, these observations highlight both the continued value of established disease-gene associations for rare disease diagnosis and the substantial opportunity within GREGoR for novel gene discovery and refinement of gene-phenotype relationships.

The storage and sharing of genetic findings through the GREGoR Data Model supports downstream phenotype-genotype matching, cross-cohort comparison, and iterative reanalysis as disease knowledge evolves. Candidate variants and genes are routinely shared through the federated GA4GH Matchmaker Exchange platform through various nodes including GeneMatcher, MyGene2, and seqr; evidence levels for gene-disease relationship assertions through the Gene Curation Coalition (GenCC^17^); classified variants are submitted to ClinVar; and ongoing efforts aim to support sharing through the emerging Variant-Level Matching network currently using seqr on AnVIL. To improve future discoverability and interoperability, GREGoR variants are also submitted to the ClinGen Allele Registry and annotated with GA4GH Variation Representation Specification (VRS) identifiers where available.

### Data Access and Analysis

Genomic data, including 8,420 short-read genome sequencing samples, along with 14 RNA sequencing samples, are available in *seqr*^18^ on the AnVIL platform for analysis. The data is annotated with relevant clinical information from the data model, including the variants reported as GREGoR findings. Within *seqr*, users can perform both case-specific and project-wide analyses, as well as utilize Matchmaker Exchange and Variant-Level Matching tools. Case status updates can be entered by the respective data submitters, and contributions from other users about potential diagnostic variants can be flagged in a case, and are then broadly available to all users with access to the case. The consortium is evaluating options for providing this *seqr* data access to anyone with permission for GREGoR data.

In addition to cloud-based analysis environments, the GREGoR Data Model has been implemented within institutional research infrastructure. At one Research Center, the canonical, version-controlled schema serves as the foundation of an internal dashboard that enforces automated validation of submitted records and supports structured experiment tracking. The system uses a modular architecture in which schema definitions, server-side validation, and client interfaces are maintained as independent components, enabling API-based interoperability with other research systems. This deployment demonstrates that the GREGoR Data Model functions not only as a data submission framework but also as an operational backbone for managing multi-omic research data. This dashboard is available as open-source software on GitHub to serve as a lightweight information management system for any laboratory looking to implement the GREGoR Data Model (https://github.com/UCI-ICTS/icts-dashboard).

## Discussion

As a multi-center rare disease research consortium, producing a coherent and shareable dataset is a foundational aspect of our work. Achieving this required the development of a shared data model that established clear conventions for representing phenotypic, familial, and multi-omic data. This common framework enabled the Consortium to harmonize heterogeneous datasets and efficiently support data quality control, submission, and release to the broader scientific community.

Although established standards support elements of clinical, phenotypic, or genomic data exchange^19–23^, to our knowledge, none were designed to serve as an organizational backbone for a multi-center rare disease genomics consortium. The GREGoR Data Model was developed through extensive collaboration and subject matter expertise to provide structured representations of phenotype, family relationships, solve status, assay-level metadata, and curated variant findings across diverse -omic assays. These representations support consistent interpretation, reanalysis, and reuse. The model does not eliminate upstream variability in data capture; heterogeneity in data entry practices and differences in clinical data availability were reflected in the completeness of optional fields, which ranged from 0.5% to 66% across tables. Continued iteration of the schema will be required to accommodate evolving technologies and clinical standards.

To support scalable access and analysis, the GREGoR Data Model is implemented on the NHGRI AnVIL platform. The model underlies data submission and validation workflows coordinated by the GREGoR DCC. Although integrated with AnVIL, the model is platform-agnostic: its schema and metadata conventions can be readily deployed in other cloud infrastructures. Its version-controlled, machine-readable schema can be deployed in other computational environments, and differences in data schemas, phenotypic conventions, or variant representation can be addressed by custom mapping or data transformation.

Beyond cloud-based data sharing, the GREGoR Data Model has also been deployed within institutional data infrastructure, where its machine-readable schema supports automated validation and standardized experiment tracking. This operational implementation demonstrates that the model functions not only as a submission and release framework, but also as a structured backbone for managing multi-omic research data.

Adoption of components of the GREGoR Data Model by complementary initiatives further illustrates its portability and interoperability. Investigators within the Undiagnosed Diseases Network (UDN) have reported that GREGoR-defined metadata fields for RNA sequencing and related assays improved annotation of uploaded files, enhancing cohort-level usability of long-read genome and transcriptome datasets. The UDN has applied these elements both retrospectively to harmonize previously submitted data and prospectively to structure ongoing submissions. Use of GREGoR Data Model components also simplified preparation of datasets for submission to dbGaP.

Similarly, investigators from Genomic Answers for Kids (GA4K)^24^ mapped metadata to the GREGoR Data Model to enable cross-cohort analysis. Harmonization of inputs across centers and sequencing technologies allowed datasets to be immediately comparable and shareable, even when derived from distinct analytic pipelines. The schema captures elements that are often difficult to reconstruct retrospectively, including solve status, affected status, proband and family relationships, HPO-coded phenotypes, consent codes, and standardized sequencing metadata (e.g., platform, chemistry, tissue), with explicit participant → analyte → experiment → aligned/variant file linkages. Alignment of these structured fields facilitated comparison of HiFi-resolved structural variants and integration of transcriptomic and methylation data across cohorts. GA4K investigators reported improved matching of controls, ancestry-aware background rate estimation, and increased statistical power without reliance on inferred diagnostic metadata.

Structured harmonization also supported broader multi-institutional collaboration. Integration of datasets from research groups studying unsolved rare disease families culminated in a Rare Disease Research Hackathon held in May 2024^25^. During this event, multidisciplinary teams applied diverse analytic approaches—including genome-wide read-depth quality control metrics, annotated structural variant frequency resources^25^, and visualization tools for high-confidence copy-number variant detection^26^—to jointly analyze genome sequencing data, including GREGoR families. Across participating cohorts, previously undetected clinically relevant variants were identified. These independent efforts illustrate how consistent metadata conventions reduce barriers to cross-cohort analysis and enable community-driven methodological innovation.

Within GREGoR, implementation of this shared framework enabled thirteen quarterly data-sharing cycles over five years, supported five public data releases, and facilitated the continued growth of the dataset into an increasingly comprehensive multi-omic resource. The Dataset now includes 12,292 participants, most of whom have short-read genomes, with substantial subsets contributing short-read RNA-seq and long-read genome data. Structured candidate variants reported across research sites provide a detailed view of prioritized genetic findings. Because the majority of participants remain unsolved (84%), future analyses have strong potential to yield new diagnoses and advance novel gene–disease discovery. Moreover, the overlapping multi-omic datasets are expected to serve as a valuable resource for methods development, benchmarking, and integrative analysis.

## Conclusions

The GREGoR Data Model and the resulting GREGoR Dataset demonstrate that a collaboratively developed, structured framework can successfully organize and sustain large-scale, multi-center rare disease genomics research. By encoding consensus metadata standards as a version-controlled, machine-readable schema with automated validation workflows, the Consortium established durable infrastructure for harmonized data submission, controlled-access release, and iterative reuse. The resulting dataset—currently comprising more than 12,000 deeply phenotyped participants with expanding multi-omic data—provides a substantial resource for gene discovery, variant interpretation, and methodological benchmarking.

More broadly, this work illustrates how intentional data structuring enables interoperability across programs and analytical platforms. As multi-omic technologies continue to evolve, progress in rare disease genomics will depend not only on data generation, but on the development of shared frameworks that maximize reuse and integration. The GREGoR Data Model offers one such framework for building interoperable, extensible human multi-omic datasets. Additional adoption across the rare disease research community will help to empower rare disease research by allowing more direct comparisons and supporting integrated analyses.

## Supporting information

Supplement 1: The GREGoR Data Model

Supplement 2: Data Model Schema

Supplement 4: Table S4. De-identified candidate genetic findings with supporting annotation

## Supplement

**Document S1**: Detailed field-level definitions for the full data model (version 1.11) .xls file

**Document S2:** Tables of the GREGoR Data Model .pdf file follows

**Document S3:**Tables S1-S3 (Solve definitions, Variant Interpretation, Completeness of Optional Tables) .pdf file follows

**Document S4:** Table S4 (Genetic Findings) .xls file

## Acknowledgments

The GREGoR Consortium is funded by the National Human Genome Research Institute of the National Institutes of Health, through the following grants: U01HG011744, U01HG011745, U01HG011755, U01HG011758, U01HG011762, and U24HG011746. The content is solely the responsibility of the authors and does not necessarily represent the official views of the National Institutes of Health.

Support for title page creation and format was provided by AuthorArranger, a tool developed at the National Cancer Institute.

## Author Contributions

BDH, MMW, CMC, MPC, ECD, SD, SMG, SNJ, CTM, AO-L, AMS, RAU, BW, members of the Data Standards and Analysis Working Group, SIB, and JXC contributed to conceptualization of this work.

BDH, MMW, CMC, SD, SMG, SNJ, CHK, LP, and members of the GREGoR Consortium contributed to data curation.

MMW, SD, and SMG contributed to formal analysis.

BDH, CMC, DEM, AO-L, GQ, MB, JB, EE, RAG, JRL, SM, SBM, TP, JP, HLR, AS, MET, EV, CW, MTW, and QY contributed to funding acquisition.

BDH, CMC, CHK, AMS, SIB, and JXC contributed to investigation.

BDH, MMW, SD, SMG, CHK, AMS, and LP contributed to methodology.

BDH, SNJ, AO-L, and the GREGoR Consortium contributed to project administration.

MPC, DEM, GQ, and the authors contributing to funding acquisition contributed resources.

MMW, SMG, SNJ, CHK, AO-L, LP, AMS, CCT, MB, JB, EE, RAG, JRL, SM, SBM, TP, JP, HLR, AS, MET, EV, CW, MTW, and QY contributed software.

BDH, LP, and AMS contributed supervision.

MMW and SMG contributed validation.

MMW and CCT contributed visualization.

BDH, MMW, CMC, SD, SMG, SNJ, CHK, LP, AO-L, AMS, CCT, RAU, BW, the GREGoR Consortium, GREGoR Consortium Data Standards and Analysis Working Group, SIB, and JXC contributed to writing of the original draft.

All authors contributed to review and editing of the submitted manuscript.

## Declaration of Interests

R. A. Gibbs and Baylor College of Medicine have equity in Codified Genomics.

D. E. Miller is on the scientific advisory board at Inso Biosciences; is engaged in research agreements with Oxford Nanopore Technologies (ONT), PacBio, Illumina, and GeneDx; has received research and travel support from ONT, PacBio, and Illumina; holds stock options in MyOme and Inso Biosciences; and is a consultant for MyOme.

S. Berger is a employee of Ambry genetics, a subsidiary of Tempus AI

S. B. Montgomery is an advisor to BridgeBio, MyOme and PhiTech.

M. J. Bamshad is on the scientific advisory board of GeneDx and has research agreements with GeneDx, Illumina, Inc., and PacBio,Inc.

H. Rehm is a shareholder and past advisory board member of Genome Medical and receives research funding from Microsoft.

J. E. Posey serves on the Scientific Advisory Board of MaddieBio.

## Notes

https://www.ncbi.nlm.nih.gov/projects/gap/cgi-bin/study.cgi?study_id=phs003047

https://anvil.terra.bio/

https://github.com/UW-GAC/gregor_data_models

## References

1. Bamshad, M.J., Nickerson, D.A., and Chong, J.X. (2019). Mendelian Gene Discovery: Fast and Furious with No End in Sight. Am. J. Hum. Genet. 105, 448–455. 10.1016/j.ajhg.2019.07.011.

2. Sullivan, J.A., Schoch, K., Spillmann, R.C., and Shashi, V. (2023). Exome/Genome Sequencing in Undiagnosed Syndromes. Annu. Rev. Med. 74, 489–502. 10.1146/annurev-med-042921-110721.

3. Baxter, S.M., Posey, J.E., Lake, N.J., Sobreira, N., Chong, J.X., Buyske, S., Blue, E.E., Chadwick, L.H., Coban-Akdemir, Z.H., Doheny, K.F., et al. (2022). Centers for Mendelian Genomics: A decade of facilitating gene discovery. Genet. Med. 24, 784–797. 10.1016/j.gim.2021.12.005.

4. Dawood, M., Heavner, B., Wheeler, M.M., Ungar, R.A., LoTempio, J., Wiel, L., Berger, S., Bernstein, J.A., Chong, J.X., Délot, E.C., et al. (2025). GREGoR: accelerating genomics for rare diseases. Nature 647, 331–342. 10.1038/s41586-025-09613-8.

5. National Institutes of Health (2025). Required Security and Operational Standards for NIH Controlled-Access Data Repositories (U.S. Department of Health and Human Services.).

6. Schatz, M.C., Philippakis, A.A., Afgan, E., Banks, E., Carey, V.J., Carroll, R.J., Culotti, A., Ellrott, K., Goecks, J., Grossman, R.L., et al. (2022). Inverting the model of genomics data sharing with the NHGRI Genomic Data Science Analysis, Visualization, and Informatics Lab-space. Cell Genomics 2, 100085. 10.1016/j.xgen.2021.100085.

7. Deciphering Developmental Disorders Study (2017). Prevalence and architecture of de novo mutations in developmental disorders. Nature 542, 433–438. 10.1038/nature21062.

8. Sharma, D.K., Solbrig, H.R., Prud’hommeaux, E., Pathak, J., and Jiang, G. (2016). Standardized Representation of Clinical Study Data Dictionaries with CIMI Archetypes. AMIA Annu. Symp. Proc. AMIA Symp. 2016, 1119–1128.

9. Vidal, M.-E., Endris, K.M., Jozashoori, S., Karim, F., and Palma, G. (2019). Semantic Data Integration of Big Biomedical Data for Supporting Personalised Medicine. In Current Trends in Semantic Web Technologies: Theory and Practice Studies in Computational Intelligence., G. Alor-Hernández, J. L. Sánchez-Cervantes, A. Rodríguez-González, and R. Valencia-García, eds. (Springer International Publishing), pp. 25–56. 10.1007/978-3-030-06149-4_2.

10. Gogarten, S., Heavner, B., Tong, C., and Stilp, A. (2025). UW-GAC/gregor_data_models: v1.10. Version v1.10 (Zenodo). 10.5281/ZENODO.14660039.

11. All of Us Research Program Investigators, Denny, J.C., Rutter, J.L., Goldstein, D.B., Philippakis, A., Smoller, J.W., Jenkins, G., and Dishman, E. (2019). The “All of Us” Research Program. N. Engl. J. Med. 381, 668–676. 10.1056/NEJMsr1809937.

12. Reich, C., Ostropolets, A., Ryan, P., Rijnbeek, P., Schuemie, M., Davydov, A., Dymshyts, D., and Hripcsak, G. (2024). OHDSI Standardized Vocabularies-a large-scale centralized reference ontology for international data harmonization. J. Am. Med. Inform. Assoc. JAMIA 31, 583–590. 10.1093/jamia/ocad247.

13. Riggs, E.R., Andersen, E.F., Cherry, A.M., Kantarci, S., Kearney, H., Patel, A., Raca, G., Ritter, D.I., South, S.T., Thorland, E.C., et al. (2020). Technical standards for the interpretation and reporting of constitutional copy-number variants: a joint consensus recommendation of the American College of Medical Genetics and Genomics (ACMG) and the Clinical Genome Resource (ClinGen). Genet. Med. Off. J. Am. Coll. Med. Genet. 22, 245–257. 10.1038/s41436-019-0686-8.

14. Strande, N.T., Riggs, E.R., Buchanan, A.H., Ceyhan-Birsoy, O., DiStefano, M., Dwight, S.S., Goldstein, J., Ghosh, R., Seifert, B.A., Sneddon, T.P., et al. (2017). Evaluating the Clinical Validity of Gene-Disease Associations: An Evidence-Based Framework Developed by the Clinical Genome Resource. Am. J. Hum. Genet. 100, 895–906. 10.1016/j.ajhg.2017.04.015.

15. Richards, S., Aziz, N., Bale, S., Bick, D., Das, S., Gastier-Foster, J., Grody, W.W., Hegde, M., Lyon, E., Spector, E., et al. (2015). Standards and guidelines for the interpretation of sequence variants: a joint consensus recommendation of the American College of Medical Genetics and Genomics and the Association for Molecular Pathology. Genet. Med. Off. J. Am. Coll. Med. Genet. 17, 405–424. 10.1038/gim.2015.30.

16. Bioc-AnVIL Team (2022). BiocAnVIL: Bioconductor’s contributions to [NHGRI’s Analysis and VIsualization Laboratory (AnVIL).

17. DiStefano, M.T., Goehringer, S., Babb, L., Alkuraya, F.S., Amberger, J., Amin, M., Austin-Tse, C., Balzotti, M., Berg, J.S., Birney, E., et al. (2022). The Gene Curation Coalition: A global effort to harmonize gene-disease evidence resources. Genet. Med. Off. J. Am. Coll. Med. Genet. 24, 1732–1742. 10.1016/j.gim.2022.04.017.

18. Pais, L.S., Snow, H., Weisburd, B., Zhang, S., Baxter, S.M., DiTroia, S., O’Heir, E., England, E., Chao, K.R., Lemire, G., et al. (2022). seqr: A web-based analysis and collaboration tool for rare disease genomics. Hum. Mutat. 43, 698–707. 10.1002/humu.24366.

19. Health Level Seven International (2024). HL7 FHIR Standard, Release 5.

20. Gargano, M.A., Matentzoglu, N., Coleman, B., Addo-Lartey, E.B., Anagnostopoulos, A.V., Anderton, J., Avillach, P., Bagley, A.M., Bakštein, E., Balhoff, J.P., et al. (2024). The Human Phenotype Ontology in 2024: phenotypes around the world. Nucleic Acids Res. 52, D1333–D1346. 10.1093/nar/gkad1005.

21. Amberger, J.S., Bocchini, C.A., Schiettecatte, F., Scott, A.F., and Hamosh, A. (2015). OMIM.org: Online Mendelian Inheritance in Man (OMIM®), an online catalog of human genes and genetic disorders. Nucleic Acids Res. 43, D789–798. 10.1093/nar/gku1205.

22. Rekerle, L., Danis, D., Rehburg, F., Graefe, A.S.L., Bily, V., Caballero-Oteyza, A., Cacheiro, P., Chimirri, L., Chong, J.X., Connelly, E., et al. (2026). GA4GH phenopacket-driven characterization of genotype-phenotype correlations in Mendelian disorders. Am. J. Hum. Genet. 113, 57–70. 10.1016/j.ajhg.2025.12.001.

23. Chang, E., and Mostafa, J. (2021). The use of SNOMED CT, 2013-2020: a literature review. J. Am. Med. Inform. Assoc. JAMIA 28, 2017–2026. 10.1093/jamia/ocab084.

24. Cohen, A.S.A., Farrow, E.G., Abdelmoity, A.T., Alaimo, J.T., Amudhavalli, S.M., Anderson, J.T., Bansal, L., Bartik, L., Baybayan, P., Belden, B., et al. (2022). Genomic answers for children: Dynamic analyses of >1000 pediatric rare disease genomes. Genet. Med. Off. J. Am. Coll. Med. Genet. 24, 1336–1348. 10.1016/j.gim.2022.02.007.

25. Lun, M.Y., Posey, J.E., Bengtsson, J.D., Du, H., Roy, R.S., Yang, L., Ochoa, S., Yuan, B., Gillentine, M., Lindstrand, A., et al. (2025). Community-Driven Copy Number Variant Discovery at Scale: Results from a Rare Disease Genomics Hackathon. Preprint at Genetic and Genomic Medicine, 10.1101/2025.08.08.25333317 10.1101/2025.08.08.25333317.

26. Du, H., Lun, M.Y., Gagarina, L., Bengtsson, J.D., Grochowski, C.M., Mehaffey, M.G., Hwang, J.P., Jhangiani, S.N., Bhamidipati, S.V., Muzny, D.M., et al. (2025). An integrated platform for concurrent structural and single-nucleotide variants improves copy-number detection and reveals pathogenic alleles in undiagnosed Mendelian families. Genome Med. 18, 16. 10.1186/s13073-025-01593-8.

