## Supplementary figures and images for "Building an Interoperable Rare Disease Multi-omic Resource: The GREGoR Data Model and Dataset"

### Supplement 2: Data Model Schema

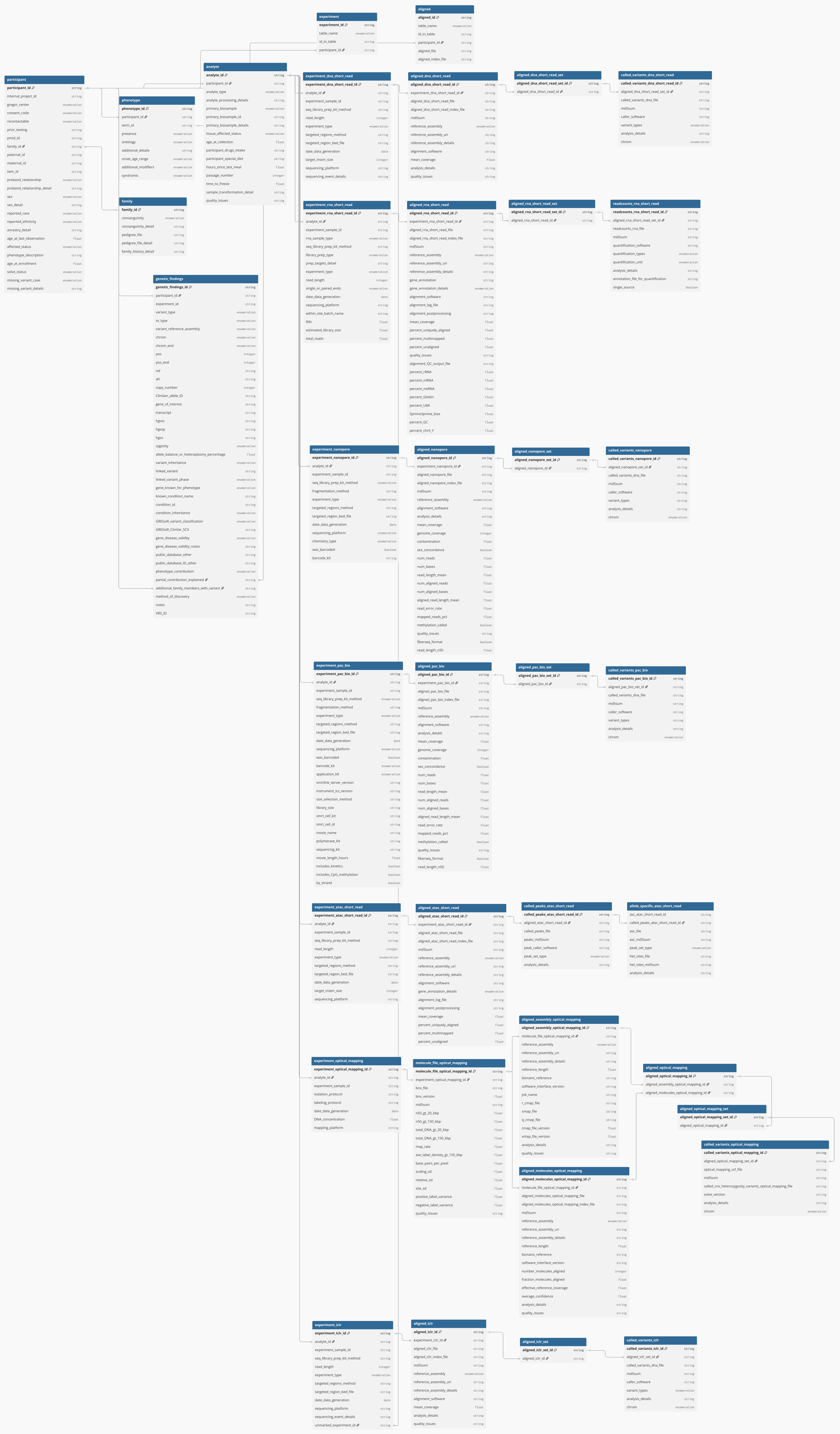


dbdiagram.io
