## Supplement 3: tables S1-S3 for "Building an Interoperable Rare Disease Multi-omic Resource: The GREGoR Data Model and Dataset"

**Table S1. Solve Status Definitions**

| Classification | Definition |
|---|---|
| **Solved** | Pathogenic or likely pathogenic variant(s) with the appropriate inheritance pattern and strong phenotype concordance in a gene with at least moderate evidence of disease association. |
| **Partially Solved** | Variant(s) explain only a subset of the observed phenotype. |
| **Probably Solved** | Variant(s) with strong but incomplete evidence, such as a high-level VUS, uncertain phase, or imperfect phenotype match, in genes with at least moderate or near-moderate gene-disease validity. |
| **Unsolved** | No variant(s) meet criteria for likely causality, including recessive cases with a missing second allele, low-confidence variants, or candidate genes with limited gene-disease evidence. |
| **Unaffected** | Individuals without the disease phenotype under investigation. |

**Table S2. Variant Interpretation by Inheritance Model**

**Dominant Model**

| Variant Evidence | Phenotype match | Classification |
|---|---|---|
| P/LP | Highly specific and consistent | Solved |
| P/LP | Consistent | Solved |
| P/LP | Imperfect | Probably Solved |
| VUS-high | Specific and consistent | Probably Solved |
| VUS-high | Imperfect | Unsolved |
| VUS-mid or lower | Any | Unsolved |

**Recessive Model**

| Variant Evidence | Phase | Phenotype match | Classification |
|---|---|---|---|

| | | | |
|---|---|---|---|
| P/LP + P/LP | Known | Specific and/or consistent | Solved |
| P/LP + P/LP | Unknown | Specific and/or consistent | Solved (if compelling) |
| P/LP + VUS-high | Any | Highly specific and/or consistent | Probably Solved |
| VUS-high + VUS-high | Known | Specific and/or consistent | Probably Solved |
| Any | Any | Inconsistent | Unsolved |
| VUS-mid or lower + VUS-mid or lower | Any | Any | Unsolved |
| Single P/LP (missing allele) | — | Any | Unsolved |

**Table S3. Completeness of Optional Fields Across Data Tables in the GREGoR Data Model**

| Table | Mean | Median | STD | Min | Max | n |
|---|---|---|---|---|---|---|
| aligned_atac_short_read | 0.67 | 1.0 | 0.5 | 0.0 | 1.0 | 9 |
| experiment_dna_short_read | 0.64 | 0.92 | 0.43 | 0.0 | 0.98 | 8 |
| experiment_rna_short_read | 0.55 | 0.63 | 0.32 | 0.0 | 1.0 | 9 |
| called_variants_nanopore | 0.52 | 0.47 | 0.46 | 0.08 | 1.0 | 3 |
| experiment_atac_short_read | 0.5 | 0.5 | 0.53 | 0.0 | 1.0 | 8 |
| readcounts_rna_short_read | 0.5 | 0.5 | 0.58 | 0.0 | 1.0 | 4 |
| genetic_findings | 0.48 | 0.55 | 0.4 | 0.0 | 1.0 | 33 |
| called_variants_pac_bio | 0.46 | 0.38 | 0.5 | 0.0 | 1.0 | 3 |
| aligned_nanopore | 0.46 | 0.49 | 0.13 | 0.0 | 0.5 | 14 |
| experiment_nanopore | 0.4 | 0.3 | 0.43 | 0.0 | 0.98 | 8 |
| called_variants_dna_short_read | 0.33 | 0.0 | 0.58 | 0.0 | 1.0 | 3 |
| participant | 0.32 | 0.3 | 0.3 | 0.0 | 0.88 | 14 |
| phenotype | 0.25 | 0.06 | 0.43 | 0.0 | 1.0 | 5 |

| | | | | | | |
|---|---|---|---|---|---|---|
| experiment_pac_bio | 0.22 | 0.0 | 0.33 | 0.0 | 1.0 | 19 |
| aligned_dna_short_read | 0.18 | 0.06 | 0.29 | 0.0 | 0.69 | 5 |
| aligned_pac_bio | 0.16 | 0.19 | 0.08 | 0.0 | 0.29 | 16 |
| aligned_rna_short_read | 0.14 | 0.0 | 0.18 | 0.0 | 0.66 | 19 |
| analyte | 0.05 | 0.0 | 0.14 | 0.0 | 0.5 | 12 |
| family | 0.03 | 0.03 | 0.04 | 0.0 | 0.07 | 4 |

n denotes the number of optional fields in the corresponding data table.
